# Tissue-specific transcriptomics uncovers novel craniofacial genes underlying jaw divergence in specialist pupfishes

**DOI:** 10.1101/2024.10.02.616385

**Authors:** M. Fernanda Palominos, Vanessa Muhl, Christopher H. Martin

## Abstract

The regulation of gene expression is one of the key evolutionary processes driving phenotypic divergence among species. Here, we investigate the tissue-specific gene expression of a non-model adaptive radiation of *Cyprinodon* pupfishes, characterized by their divergent dietary niches and exceptionally fast rates of craniofacial evolution. By comparing tissue-specific gene expression in the most morphologically divergent skeletal structure, the oral jaws, with the relatively morphologically conserved caudal tail region, we identified genes that were differentially expressed exclusively in the developing jaws of each of the three trophic specialists at hatching (8 dpf) and not in any other species. We then assessed their overlap (as transcriptionally-regulated genes) with adaptive regulatory variants identified in previous genomic studies. Our analysis identified *pycr3* and *atp8a1* as the most promising for craniofacial evolution in the scale-eaters, both genes with no known previous craniofacial function. We functionally confirmed the craniofacial expression of these genes by *in situ* mRNA hybridization chain reaction and demonstrated their species-specific expression in branchial and muscle tissues between sister species of this young radiation. Our work underscores the power of integrating tissue-specific transcriptomics with speciation genomics to identify novel craniofacial candidate genes controlling divergent morphogenesis in a natural ‘evolutionary mutant’ system.

## Introduction

Adaptive radiations are especially informative for understanding the genetic bases of phenotypic evolution in vertebrates (Gillespie et al. 2020; Martin and Richards 2019; Powder and Albertson 2016). Young adaptive radiations, in particular, provide a framework for studying the genetic variation that leads to phenotypic transitions in systems with high rates of hybridization and minimal extinction (Schluter 2000). A classic example of this is the Galápagos radiation of Darwin’s finches, in which *bmp4* (*bone morphogenetic protein 4*) levels and timing of expression vary among finches during craniofacial development (Abzhanov et al. 2004) and variants in the coding and non-coding regions of *alx1* (*Alx homeobox 1*) control differences in size and bluntness of the beak (Lamichhaney et al. 2015; Rubin et al. 2022). Studies on the immense craniofacial diversity within adaptive radiations of fishes have shown that the co-option of known craniofacial genes leads to the expansion or reduction of their expression field allowing for the development of divergent craniofacial traits between species and populations. For example, a genetic substitution in the *bmp6* (*bone morphogenetic protein 6*) enhancer region is able to control the increased tooth number observed in derived freshwater three-spined sticklebacks and not in the marine populations (Cleves et al. 2018; Stepaniak, Square, and Miller 2021). Similarly, craniofacial evolution of cichlids in Africa’s Great Lakes has been linked to changes in expression and function of several other previously known craniofacial genes, such as *bmp4* (Albertson et al. 2005; Powder et al. 2014), *fgf* (Albertson et al. 2018), *lbh* (Powder et al. 2014), *ptch1* (Roberts et al. 2011; Hu and Albertson 2014; DeLorenzo et al. 2022), *sox9b* (Matthews and Albertson 2017), among others (Streelman et al. 2003; Conith and Albertson 2021).

Despite advances in understanding how well-known craniofacial genes contribute to the evolution of divergent morphologies among radiating species, the potential of adaptive radiations to identify novel craniofacial gene function in vertebrates has been largely overlooked (but see discovery of facial bone fragmentation in the Mexican cavefish (Powers et al. 2023)). Discovering new gene regulatory networks involved in craniofacial development can provide valuable insights into subtle phenotypic changes observed in natural systems, which potentially may linked to rare and small-effect human craniofacial genetic syndromes (Powder and Albertson 2016; Albertson et al. 2009; Streelman, Peichel, and Parichy 2007).

The nascent *Cyprinodon* adaptive radiation is comprised of four pupfish species occuring in sympatry in the hypersaline lakes of San Salvador Island (SSI), Bahamas (Martin and Wainwright 2013a, 2013b). This endemic radiation comprises three trophic specialists: the molluscivore pupfish (*C. brontotheroides*), with a novel nasal and maxillary skeletal protrusion; the scale-eating pupfish (*C. desquamator*) with greatly enlarged oral jaws and adductor mandibulae muscles (Hernandez et al. 2018); and a recently discovered intermediate scale-eater (*C.* sp. ‘wide-mouth’), which possess the widest jaws and intermediate jaw lengths (Richards and Martin 2022). All trophic specialists are derived from a Caribbean-wide generalist (*C. variegatus*) also present in SSI’s lakes. Originating approximately 10,000 years ago, the SSI pupfish radiation rapidly evolved divergent and unique craniofacial phenotypes despite minimal genetic divergence (Martin and Wainwright 2011; Martin, Erickson, and Miller 2017; McGirr and Martin 2017, 2021) and ongoing gene flow (Richards and Martin 2017; Richards et al. 2021; Patton et al. 2022). Notably, SSI pupfishes are highly amenable to laboratory rearing, imaging, and developmental biology studies (Palominos et al. 2023), making them a unique model to study the genetic bases of craniofacial diversification. Previously, we have demonstrated its advantages to discover a novel function for *galr2* (*galanin receptor 2*) in craniofacial development in lower oral jaw elongation in the scale-eater pupfish, highlighting the potential of this system to reveal novel genetic function in vertebrate craniofacial adaptation and disease.

Gene expression regulation is arguably one of the main mechanisms driving phenotypic diversity among closely related species. In SSI, the specialized craniofacial morphologies of each pupfish species are apparent as early as hatching (E. Lencer and McCune 2020; E. S. Lencer, Riccio, and McCune 2016; Holtmeier 2001; Palominos et al. 2023), and genetically determined through multiple generations of laboratory rearing in a common garden. Previous transcriptomic studies in this system identified differentially expressed genes (DEG) between species in known canonical key craniofacial developmental pathways such as IgF and Wnt (E. S. Lencer et al. 2017) and cell cycle regulation (E. Lencer and McCune 2020). Earlier work from our lab has linked genetic variants with gene expression changes in whole embryos and larvae and identified two novel craniofacial candidate genes *dync2li1* and *pycr3*, which were differentially expressed throughout embryonic and larval development (McGirr and Martin 2021). However, differentially DEGs between species have not yet been studied in a tissue-specific manner, nor have included outgroup pupfish species outside SSI to compare in this context. In this study, we applied high-depth tissue-specific RNA sequencing to compare the oral jaws -exhibiting the highest morphological divergence within the radiation-to the caudal region (or ‘tail’), which displays minimal divergence between species. We compared four pupfish species within this radiation (generalist, molluscivore, scale-eater, and ‘wide-mouth’) and two generalist outgroup species to identify and validate novel jaw-specific DEGs at hatching associated with fixed genomic variants unique to SSI trophic specialists.

## Methods

### Study system and pupfish husbandry

*Cyprinodon* species were maintained in the lab for at least two generations in small breeding groups. Two lake populations of scale-eaters, two lake populations of molluscivores, one population of intermediate scale-eater, and four populations of generalists were reared in the lab for multiple generations and sampled for tissue-specific gene expression at 8 dpf. From San Salvador Island, *C. variegatus* and *C. brontotheroides* were collected from Osprey Lake and Crescent Pond in 2018, while *C. desquamator* individuals were collected from Little Lake in 2014 (which is connected to Osprey Lake through the large interior Great Lake) and Crescent Pond in 2018. A fourth sympatric and newly described intermediate scale-eater pupfish from San Salvador Island, *C.* sp. ‘wide-mouth’, was collected from Osprey Lake in 2018 (Richards and Martin 2022). We also included two outgroup generalist populations: *C. variegatus* from North Carolina in 2018, and the more distant outgroup *C. fontinalis* from Ojo de Carbonera in the Chihuahuan Desert in Mexico obtained from the American Killifish Association captive breeding program. These outgroups represent approximately 10,000 and 30,000 years of divergence from the San Salvador Island species flock, respectively (Martin et al. 2016; Martin 2016; Tian et al. 2022).

### Tissue-specific sample collection

Embryos were collected within 24 hours after natural fertilization in mixed breeding tanks and moved to a Petri dish for constant incubation at 27 °C in 2-5 ppt salinity dechlorinated water with a diluted addition of methylene blue and gentamycin to prevent fungus. Newly hatched larvae (8 dpf) from all species and populations were collected in 1 ml sterile microcentrifuge tubes filled with the RNA stabilizer RNA*later* (Thermo Fisher Scientific, Inc.), and stored for 24 hours at 4 °C, followed by long-term storage at -20 °C. Under a stereomicroscope we dissected the oral jaws and associated craniofacial connective tissues, including the dentary, angular articular, maxilla, premaxilla, palatine, and pharyngeal apparatus from each embryo (Fig. 1). Five to six embryos were pooled per biological replicate to obtain sufficient extracted RNA for sequencing libraries. We similarly dissected the caudal region from each embryo starting from the last 6 myotomes from posterior to anterior, with the caudal fin included (Fig. 1). Petri dishes, spring scissors, and forceps used for dissection were washed with RNAse AWAY (Thermo Fisher, Inc.) before use. The dissections were carried out in a sterile 30 mm plastic Petri dish filled with RNA*later* using 2mm fine Spring scissors and fine Dumont forceps (Fine Science Tools, Inc.).

**Figure 1.**
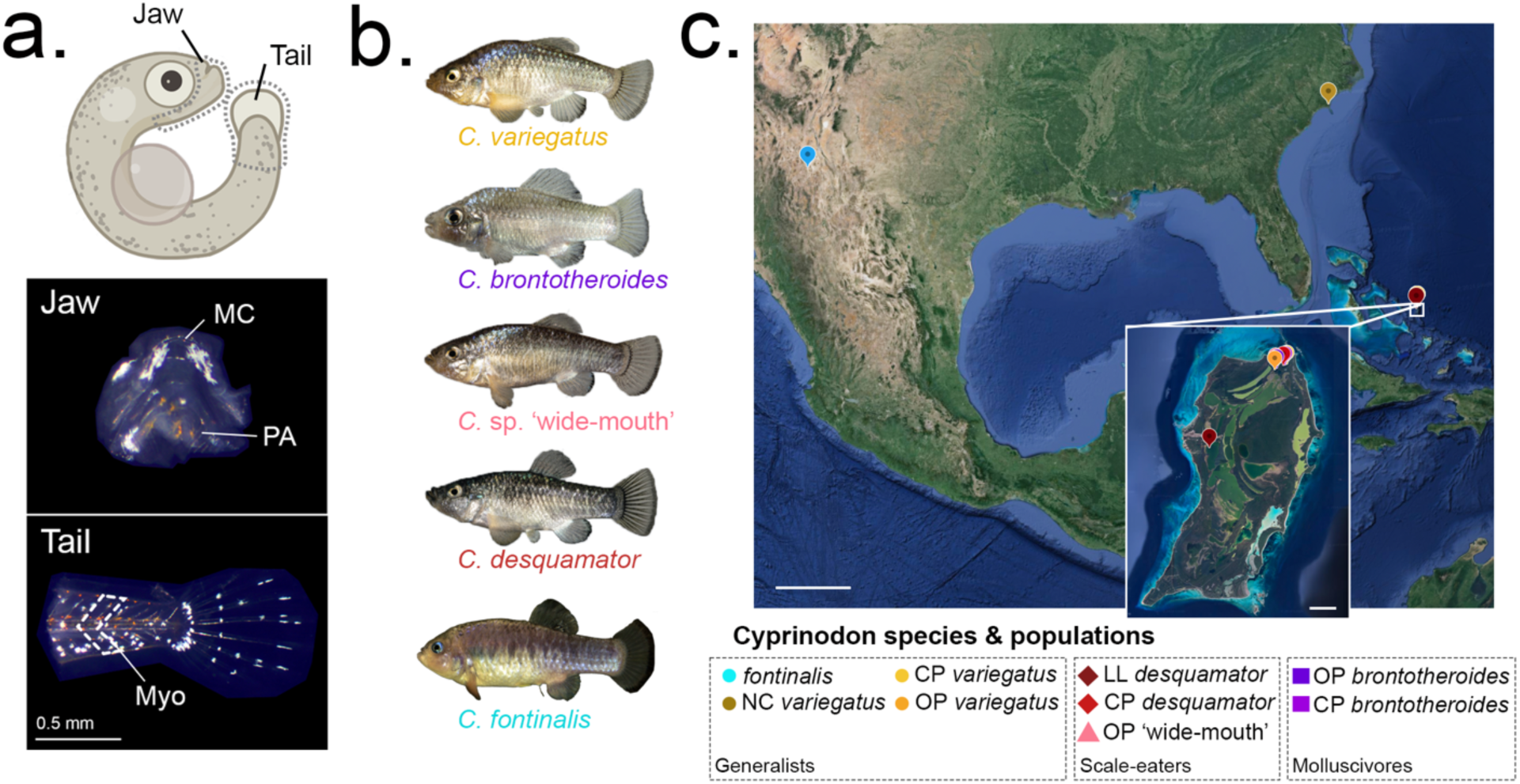
Tissue- and species-specific RNA sequencing of the San Salvador Island adaptive radiation and outgroup species at hatching. a) RNA isolation from ‘jaw’ samples included the dissection of the nasal, maxilla, palatine, premaxilla, Meckel’s cartilage (lower larval oral jaw), and the branchial apparatus (from the 1^st^ to 4^th^ pharyngeal arch). ‘Tail’ samples were dissected from the 6^th^ myotome to the caudal fin. Hatched larvae cartoon adapted from Biorender. b-c) Tissue-specific RNA was isolated from 6 lab-reared pupfish species, across 9 different populations. Inset: San Salvador Island, scale: 2 km. Larger scale: 500 km. Satellite images from Google Earth. NC: North Carolina, CP: Crescent Pond, OP: Osprey Lake, LL: Little Lake.

### RNA isolation, sequencing, and alignment

The RNA biological replicates were from “jaws” and “tails”. The tissue-specific RNA from each of the biological replicates (for each species, population, and tissue) was extracted using Monarch® Total RNA Miniprep Kit (New England Biolabs), following the manufacturer’s instructions. We validated RNA integrity using an Agilent Bioanalyzer and only samples with a RIN score > 8 were used for library preparation. The poly-A enriched libraries and sequencing were carried out by the DNA Technologies and Expression Analysis Core at the UC Davis Genome Center, supported by NIH Shared Instrumentation Grant 1S10OD010786-01. All sequenced samples are listed in Table S1.

We used 150 bp paired-end sequencing on two lanes of Illumina NovaSeq 6000 S4, resulting in 7.6 billion raw reads. We filtered raw reads using Trim Galore (v. 4.4, Babraham Bioinformatics) to remove Illumina adaptors and low-quality reads (mean Phred score < 20). We mapped reads of all species to the *Cyprinodon variegatus* reference genome (C_variegatus-1.0, GCA_000732505.1) using the RNA-seq aligner STAR (v. 2.7.10b, parameters = - outFilterMultimapNmax 1, -outFilterMismatchNmax 3). After alignment, reads were counted using HTseq (Anders, Pyl, and Huber 2015). Mapping and read quality were assessed using MultiQC (Ewels et al. 2016).

### Differential expression analyses

We normalized counts and performed differential gene expression analysis using DESeq2 (v.1.40.2; (Love, Huber, and Anders 2014) in R Studio (Racine 2012). Genes with a mean read count greater than 10 across samples were retained for analyses resulting in a set of 16,356 and 14,338 genes that were expressed in our jaw and tail datasets, respectively. We compared gene expression a) between tissues overall, b) between each species against all other species in our dataset, and c) between tissues within each species. SSI species were pooled together across both lake populations unless otherwise stated. Differential expression between comparisons was determined using Wald tests of the normalized posterior log fold change estimates and corrected for multiple testing using the Benjamini-Hochberg method with a false discovery rate (FDR) of 0.01. To categorize the potential function of craniofacial-exclusive DEGs in each pupfish species comparison, we used gene ontology enrichment analyses using ShinyGo (Ge, Jung, and Yao 2020) with the available annotations for *C. variegatus*, and via PANTHER with annotations for zebrafish (PantherRef).

### In-situ mRNA hybridization

We used chromogenic in-situ hybridization for *pycr3* in cryosections using RNAscope 2.5 (Advanced Cell Diagnostics, Inc.) performed by UNC Translational Pathology and Histopathology Core Services. 8 dpf formalin-fixed and paraffin-embedded embryos were sectioned and hybridized to a custom pupfish RNAscope probe on the Leica Biosystems’ Bond RX fully automated system with the RNAscope 2.5 LS BROWN assay kit. Fluorescent in-situ hybridizations for *atp8a1* and *tpm3b* were performed for all species at 8 dpf using HCR custom probes from Molecular Instruments, Inc. following the (Palominos et al. 2023) protocol. Samples were imaged using a Zeiss LSM880 (inverted) confocal microscope at the CNR Biological Imaging Facility at the University of California, Berkeley following Palominos et al., 2023.

## Results

### Gene expression divergence across tissues, species, and populations

To investigate the tissue- and species-specific transcriptional divergence underlying the previously reported craniofacial differences in SSI pupfishes (Palominos et al. 2023; E. Lencer and McCune 2020; E. S. Lencer, Riccio, and McCune 2016; Holtmeier 2001), we conducted RNA sequencing on isolated jaws and tails (Fig. 1a). We sampled two populations per generalist and specialist on SSI, one outgroup generalist population from North Carolina (NC), and a distant outgroup *Cyprinodon* species from the Chihuahua desert (Fig. 1b-c). The sampling design allowed to capture distant transcriptional variation associated with craniofacial evolution and dietary specialization.

Principal component analysis (PCA) revealed that tissue type and species identity were the primary drivers of transcriptional divergences, accounting for 70% and 9.5% of the variation, respectively, across 7.6 billion mapped reads (Fig. 2a-b). The PCA results demonstrated clear clustering of replicates according to sample type, species, and tissues. However, when comparing jaw tissues, SSI trophic specialists formed a distinct cluster, separated from all the generalists (*C. variegatus* from SSI and NC, plus *C. fontinalis*; Fig. 2c-e). Within this specialist cluster, scale-eaters from Crescent Pond (Fig. 2, bright red diamonds) formed a transcriptionally distinct cluster from all other jaw tissues (Fig. 2c, d) which was not different for tail tissues when focusing on SSI exclusively (Fig. 2e).

**Figure 2.**
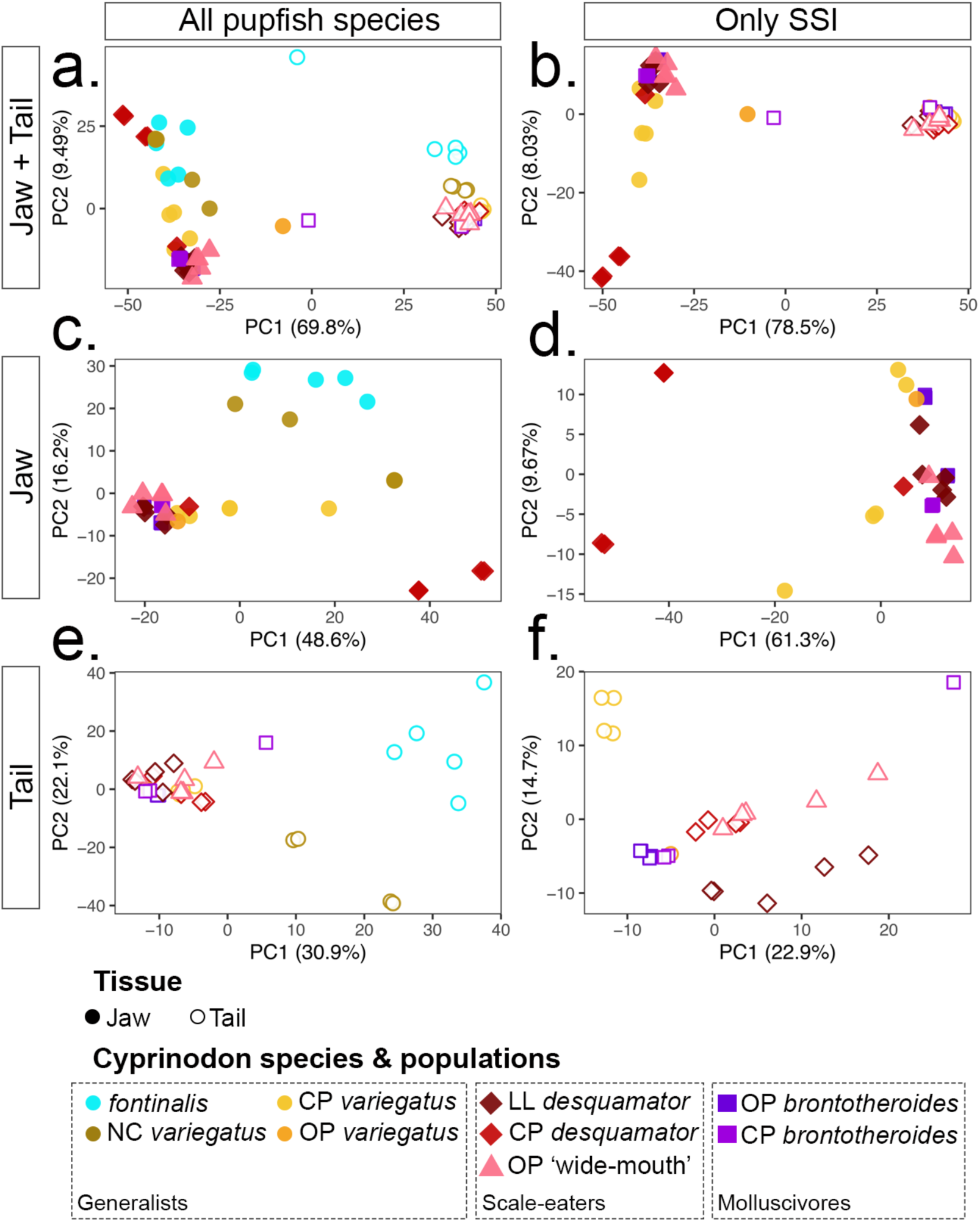
Principal component analyses of tissue-specific transcriptional variation across pupfish species and tissue-type. a-b) jaw and tail samples, c-d) only jaw tissues, and e-f) only tail tissues. First column: all 6 pupfish species, second column: SSI only: the 4 species of pupfish from San Salvador Island (*C. variegatus*, *C. desquamator*, *C. brontotheroides*, and C. sp. ‘wide-mouth’). Filled symbols: jaw tissue, open symbols: tail tissue. Pupfish populations: NC: North Carolina, CP: Crescent Pond, OP: Osprey Lake, LL: Little Lake.

### High proportions of differentially expressed genes are unique to the jaw tissues of trophic specialists relative to tail tissues

We employed DEseq2 (Love, Huber, and Anders 2014) to quantify tissue-specific differential gene expression in each specialist for each tissue type. Notably, both the molluscivore, *C. brontotheroide*s, and the intermediate scale-eater, *C.* ‘wide-mouth’, showed a high proportion of DEGs, compared to all other jaw samples with 84.4% and 94.8% of them being jaw-exclusive, respectively (Fig. 3b, purple and pink, respectively). In *C. brontotheroide*s, this corresponded to 1,656 jaw-exclusive DEGs (Fig. 3b: purple), while ‘wide-mouth’ showed 3,552 DEGs (Fig. 3b: pink) to be jaw-exclusive and differentially expressed (DE) between pupfish species. In both species, the percentage of DEGs shared between jaw and tail tissues (6.8% and 3.7%, respectively) and tail-exclusive DEGs (8.8% and 1.5%, respectively) was less than 10%. This likely reflects the substantial transcriptional divergence of the craniofacial tissue between all pupfish species in our dataset.

**Figure 3.**
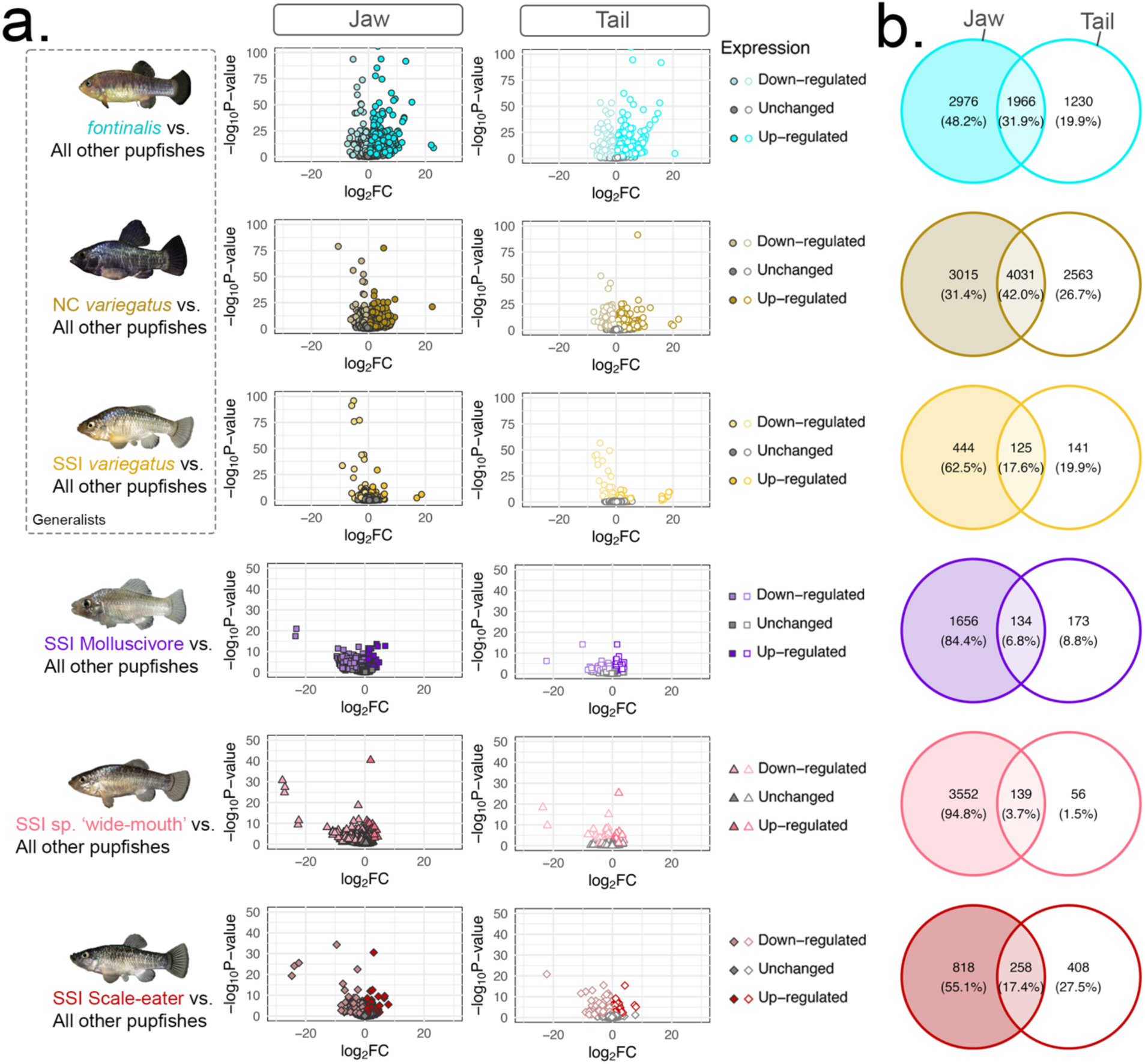
Differential gene expression in each species for each tissue type. Each row represents DEGs unique to each species relative to all species in our dataset. SSI lake populations were pooled for each comparison. Generalist species are grouped inside the dashed rectangle. a) Volcano plots showing up and downregulated genes between species, in each tissue. Filled symbols represent jaws (left panels) and open symbols represent tails (right panel). b) The number of tissue- and species-species DEGs for each comparison. Filled circle: jaw-tissue DEGs, unfilled circle: tail-tissue DEGs, and overlapping DEGs.

In contrast, fewer jaw-specific DEGs were observed in generalist pupfishes. In *C. fontinalis*, 48.2% of the DEGs were jaw-exclusive, while in the North Carolina generalist, a 31.4%, (Fig. 3b: turquoise and dark yellow, respectively), consistent with their highly similar generalist craniofacial morphology but much greater genetic divergence. When comparing the SSI generalist to all other pupfishes, we found 6.7 times less DEGs in both jaw and tail tissues (444 jaw-exclusive DEGs, and 141 tail-exclusive DEGs; Fig. 3b, light yellow).

In the scale-eater *C. desquamator*’s developing craniofacial tissues, we found only 818 DEGs (55.1%) between the jaws of scale-eaters and all other pupfishes (Fig. 3b, red). To investigate whether this smaller percentage of craniofacial-exclusive transcripts in the scale-eaters depends on the transcriptional variance observed between populations (Fig. 2), we compared each of the scale-eater populations separately (Crescent Pond and Little Lake) to all other pupfishes.

In scale-eaters from Crescent Pond, 77.2% of DEGs were jaw-exclusive compared to all other species (Fig. S1, bright red), whereas scale-eaters from Little Lake showed 53.8% jaw-exclusive DEGs (Fig. S1, dark red), with a notable increase in tail-exclusive DEGs from 12.7% to 40.4%. This variation in differential gene expression between *desquamator* populations is consistent with their genetic divergence between these populations, which is comparable to the divergence among SSI species (Turner et al. 2008; McGirr and Martin 2017) despite the populations being morphologically indistinguishable from one another in their craniofacial traits (Martin 2016). To identify the genetic factors controlling craniofacial divergence among pupfish species, we subsequently analyzed pooled lake populations for each trophic specialist.

### Enriched gene ontology terms differ between shared and unique craniofacial DEGs

Gene ontology (GO) analysis revealed distinct enrichment patterns among shared and unique craniofacial DEGs across specialist species. We identified 71 shared DEGs in jaw tissues among specialists, whereas 550, 1096, and 2925 DEGs were specific to *desquamator*, *brontotheroides*, and the ‘wide-mouth’, respectively (Fig. 4a). Shared craniofacial DEGs among specialists (Fig. 4b-c; Table S4) were significantly enriched for terms related to ‘Detection of external stimulus’, ‘Sensory perception of light stimulus’, and ‘Response to stimulus’. *C*. *brontotheroides*-specific jaw transcripts were enriched for ‘Transport’, ‘Monosaccharide metabolic process’, ‘Glucose metabolic process’, ‘Interspecies interaction between organisms’, and ‘Reproduction’ (Fig. 4d-e). In contrast, *desquamator-*specific DEGs were enriched for ‘Hemostasis’, ‘Blood coagulation’, ‘Immune system process’, and ‘Locomotion’ (Fig. 4f-g). Finally, *C.* sp. ‘wide-mouth’ scale-eaters exhibited enrichment for terms related to for ‘Neuron projection development’ and ‘Generation of neurons’ (Fig. 4h).

**Figure 4.**
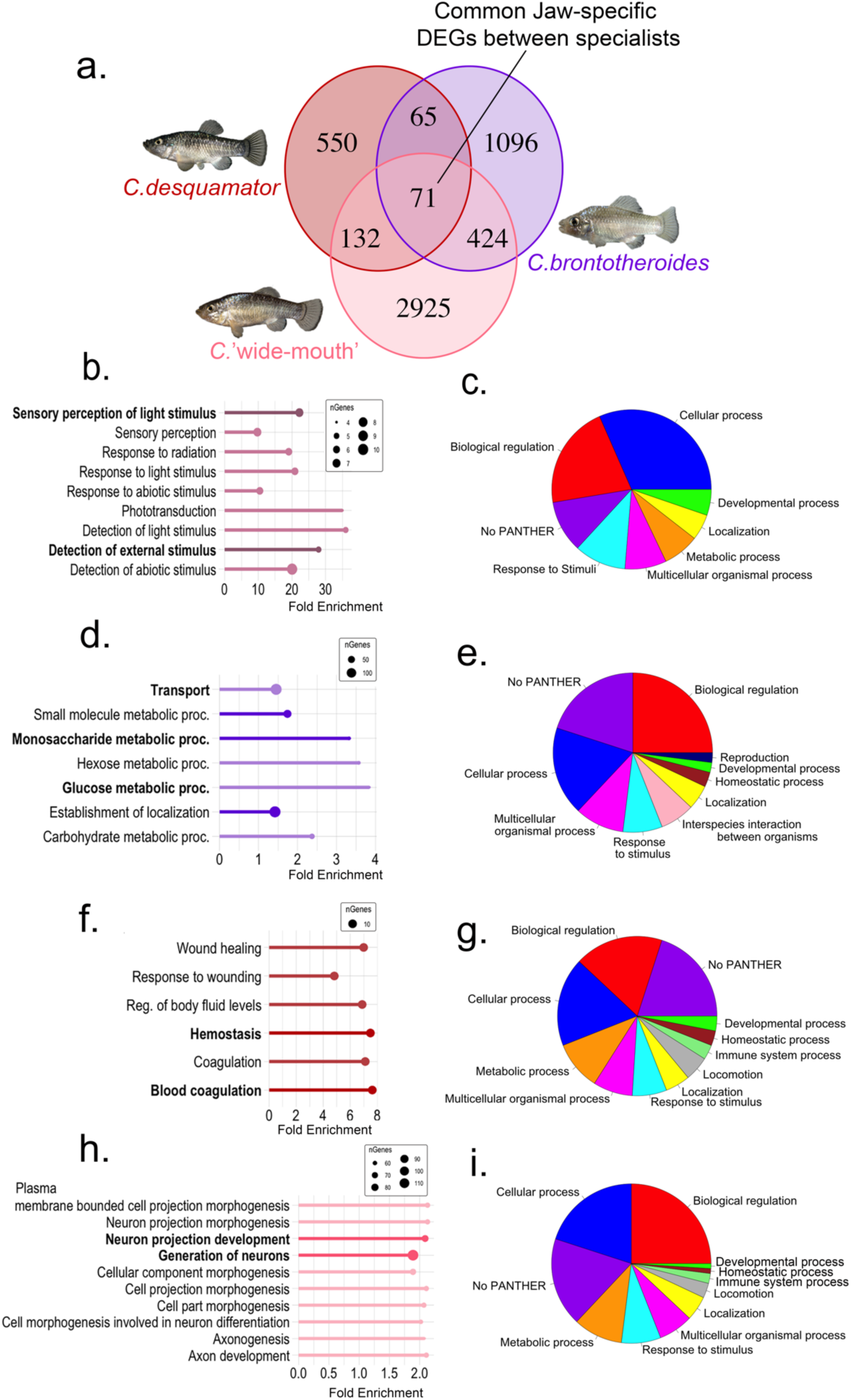
Gene ontology enrichment for shared and specialist-specific DEGs in jaw tissues. Gene ontology enrichment terms obtained using b, d, f, h) ShinyGo 0.8 and c, e, g, i) PANTHER for b, c) shared DEGs and specialists-specific DEGs in d, e) *C. brontotheroides*, f, g) *C. desquamator*, and h, i) *C. sp.* ‘wide-mouth’, relative to all other samples. Bold terms in b, d, f, h) have an FDR < 0.01.

### Convergence of adaptive alleles with species-specific DEGs in jaw tissues reveals novel craniofacial gene candidates

Previous genomic scans of 202 Caribbean pupfish genomes identified 3,436 candidate adaptive single nucleotide polymorphism (SNPs) in scale-eaters and 1,490 in molluscivores, characterized by high genetic differentiation (Fst > 0.95) between specialists. These SNPs were located within regions showing evidence of hard selective sweeps, as supported by both linkage disequilibrium and site frequency spectrum-based statistics, indicating a strong selection at these loci (Richards et al. 2021). Remarkably, 89% and 99% of the candidate alleles in molluscivores and scale-eaters, respectively, were found within 20 kb of a gene, suggesting a regulatory role in craniofacial gene expression during development. Additionally, 51 candidate adaptive alleles were identified the newly discovered intermediate scale-eater, *C.* sp. ‘wide-mouth’, using a similar pipeline (Richards and Martin 2022).

We hypothesized that by intersecting our specialist-specific DEGs exclusive to jaw tissues with these specialists’ candidate adaptive alleles would reveal novel genes where regulatory region divergence corresponds to gene expression difference between each specialist and the othet pupfish species. We found that the intersection between candidate adaptive alleles and species-specific jaw-exclusive DEGs pointed to only a few candidate genes with unknown craniofacial expression or function. We found *coq7, gga1,* and *mylipa* to be the only *brontotheroides*-specific jaw-exclusive DEGs containing adaptive alleles within 20 kb in our dataset (Fig. 5a). *Pycr3, slc51a, bri3bp, vgll3,* and *mag* were the only *desquamator*-specific jaw-exclusive DEGs containing neighboring adaptive alleles (Fig. 5b). Finally, we found only a single gene, *atp8a1,* in the *‘*wide-mouth’-specific jaw-exclusive DEGs, which contained only one single candidate adaptive allele in its regulatory region (Fig. 5c). To determine the gene expression variation of these novel craniofacial candidate genes among pupfishes we quantified their mean normalized read counts across samples.

**Figure 5.**
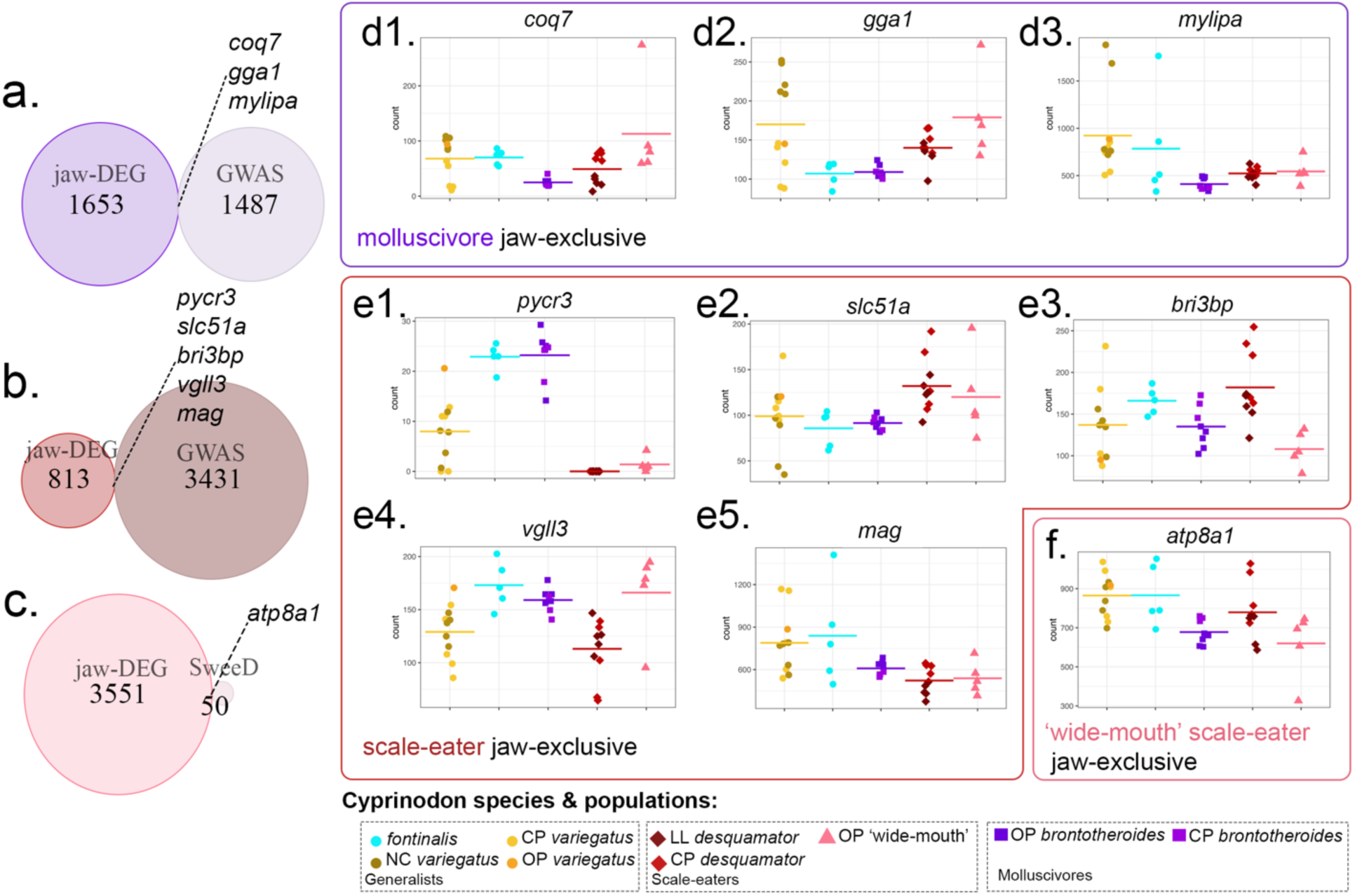
Species-specific DEGs observed only in jaw tissues overlap with few adaptive loci in each trophic specialist. Intersection between species-specific jaw-exclusive DEGs (left circles; from Fig. 3) and genes near candidate adaptive alleles (right circles; identified from genome-wide scans for highly differentiated SNPs within hard selective sweeps in Richards et al. 2021, 2022) in a) *C. brontotheroides*, b) *C. desquamator*, and c) *C.* sp. ‘wide-mouth’. Circles are scaled by the number of genes they contain. Dashed lines indicate the number of species-specific DEGs observed only in jaw tissues overlapping with genes within 20 kb of a candidate adaptive locus in each specialist. Counts across species and populations for the intersected genes identified in d1- d3) *C. brontotheroides*, e1-e5) *C. desquamator*, and f) *C.* sp. ‘wide-mouth’.

*Coq7* (*coenzyme Q7, hydroxylase*) displayed the highest log_2_-fold change and the lowest adjusted *P-*value (Table S2) in the molluscivore pupfish (Fig. 5, d1), while *gga1* (*golgi-associated, gamma adaptin ear containing, ARF binding protein 1*), interestingly, showed a decreased normalized read count only across SSI species (Fig. 5, d2,). In *C. desquamator*, *pycr3* (*pyrroline- 5-carboxylate reductase 3*) showed nearly 7-fold lower expression in craniofacial tissues compared to all other species (Table S3, Fig. 5e), with reduced expression observed in both scale-eaters, *C. desquamator* and *C. ‘*wide-mouth’. Additionally, *slc51a* (*solute carrier family 51 member A*) and *bri3bp* (*BRI3 binding protein*) showed higher expression levels in the craniofacial tissue of *desquamator* than in all other pupfishes (Fig. 5, e2, e3). Conversely, *mylipa* (myosin regulatory light chain interacting protein) and *mag* (*myelin-associated glycoprotein*) exhibited lower read count in all three specialists compared than all the generalist populations (Fig. 5, d3, e5). In the intermediate scale-eater, *atp8a1* (*ATPase phospholipid transporting 8A1*) -the only jaw-exclusive DEG associated with a nearby candidate adaptive allele- showed reduced expression relative to the other species (Fig. 5, f; Table S4). Overall, 7 out of 9 specialist-specific, jaw-exclusive DEGs (*coq7, gga1, mylipa, pycr3, vgll3, mag,* and *atp8a1*) exhibited a negative log2-fold change in gene expression relative to all other species, suggesting a species-specific transcriptional downregulation mechanism for directing divergent craniofacial development.

### Divergent expression of *pycr3* and *atp8a1* in craniofacial tissues between scale-eaters and generalist pupfishes

#### Pycr3

Among species-specific DEGs unique to craniofacial tissues near candidate adaptive loci (Fig. 5), *pycr3* was also previously identified as one of the top DEGs differentiating scale-eaters from molluscivores in whole embryos transcriptomes (McGirr and Martin 2021), which is coincident with the identification of eight different alleles within the 20 kb cis-regulatory region adjacent to *pycr3*. One of these candidate adaptive variants was an A > C transversion fixed in Crescent Pond and Osprey Lake scale-eaters, located 1,808 bp downstream of *pycr3*, and which disrupts a putative *mfz1* (*murine frizzled-*1) transcription factor binding site (McGirr and Martin 2021). Members of pyrroline-5-carboxylate reductase family, including *pycr3*, catalyze the final steps in the biosynthesis of proline with demonstrated roles in the progression of several cancer types (Burke et al. 2020; Zhuang, Huang, and Liu 2024; Bogner, Stiers, and Tanner 2021). However, their role in craniofacial development or evolution remains unknown.

To investigate the potential species-specific differential expression of *pycr3* between craniofacial tissues of the specialists, we conducted RNAscope *in-situ* mRNA hybridization assays at hatching. All pupfish species and populations analyzed showed high overall *pycr3* expression in the brain (Fig. 6, b-f), consistent with previous literature in other vertebrate models (Escande-Beillard et al. 2020; Bogner, Stiers, and Tanner 2021). However, and in line with the negligible tissue-specific expression levels of *pycr3* observed in scale-eaters (Fig. 5, e1), *pycr3* expression was absent *in situ* in the posterior skeletal elements of the jaw (gill arches 2 to 5) in scale-eaters (Fig. 6b-b1, n = 2). In contrast, *pycr3* was abundantly expressed across all five gill arches in the molluscivore hatchlings (Fig. 6a-a1, c, *n* = 2). Expression of *pycr3* in the premaxilla and future dentary bone in 1 out of the 2 individuals we analyzed (Fig. 6d-d1, n=2).

**Figure 6.**
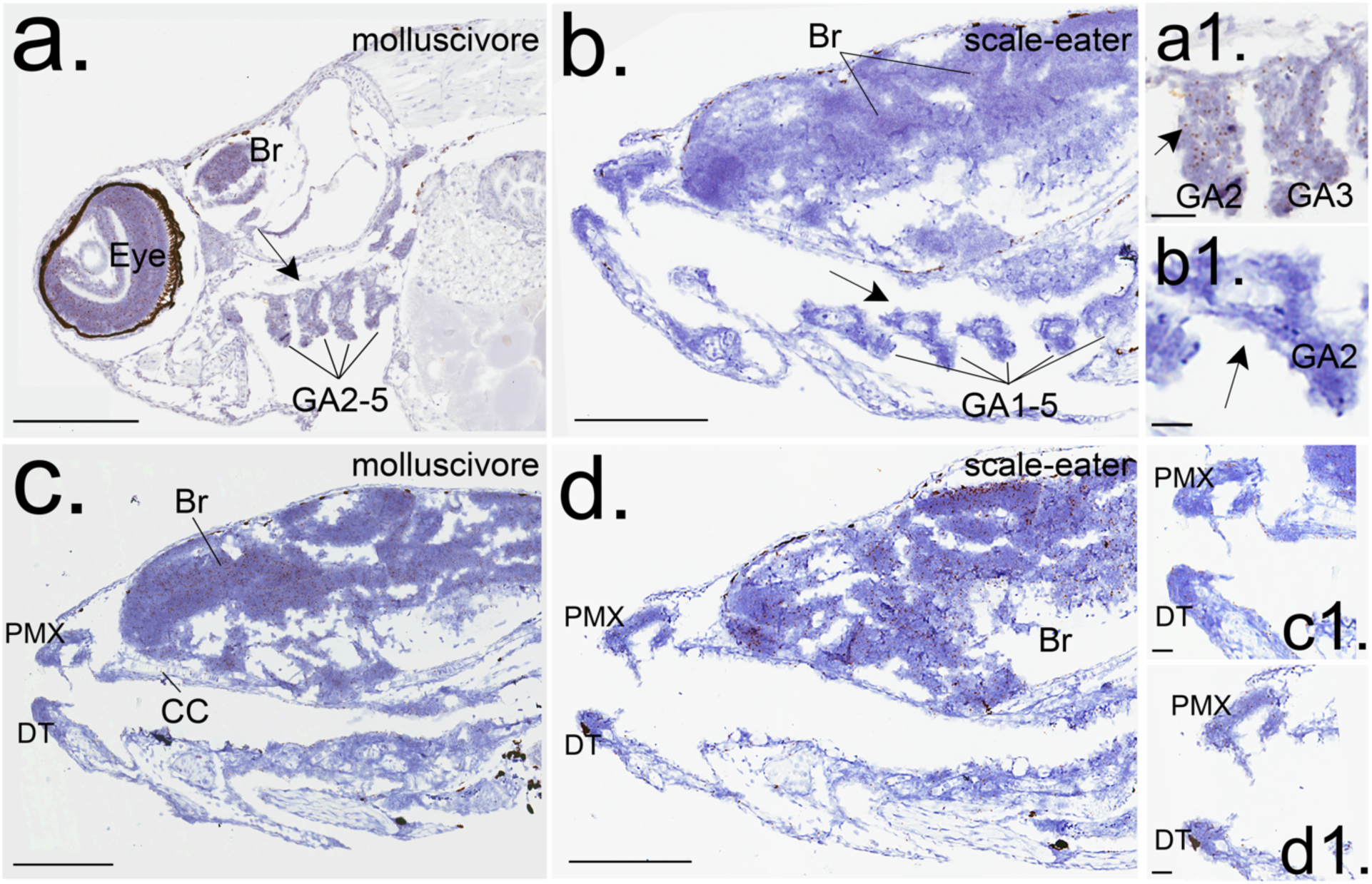
Differential *pycr3* expression in the gill arches at hatching between trophic specialists. RNAscope 2.5 in-situ hybridization for *pycr3* (dark brown spots) at hatching: b, d) *C. desquamator*; a, c) *C. brontotheroides*. GA: gill arches, Br: brain, PMX: premaxilla, DT: dentary, CC: chondrocranium. Arrows points to the difference in *pycr3* expression between specialists in the a1, b1) second gill arch (GA2). Scale bars: a-d) 200 µm, a1-d1) 25 µm.

#### Atp8a1

Among all the species-specific DEGs unique to craniofacial tissues near candidate adaptive loci (Fig. 5), *atp8a1* stood out as the only gene DE between the two scale-eaters, *C. desquamator* and *C.* ‘wide-mouth’, with a single fixed allele in scale-eaters compared to molluscivores on SSI, observed as standing genetic variation on generalist pupfish populations on other Caribbean islands (Richards et al. 2021; Richards and Martin 2022). Unlike *pycr3,* the adaptive allele near *atp8a1* was not detected in molluscivores (Richards and Martin 2022). With ubiquitous expression in humans, *atp8a1’s* function is to catalyze the hydrolysis of ATP coupled to the transport of aminophospholipids, such as phosphatidylserine (PS), from the outer to the inner membrane, controlling phospholipid asymmetry across the membrane (Hiraizumi et al. 2019).

To investigate the spatial expression of *atp8a1* in craniofacial tissues, we performed fluorescent hybridization chain reaction (HCR) for *atp8a1* and *tpm3b* (*tropomyosin 3b*), a marker for cranial muscle (Palominos et al. 2023). At hatching, *atp8a1* was broadly expressed in craniofacial tissues of generalists (Fig. 7a-a2), whereas in *desquamator*, *atp8a1* expression was mainly restricted to the intermandibular anterior (IMA) and the adductor mandibulae (AM) muscles, with significant colocalization with *tpm3b* (Fig. 7b-c).

**Figure 7.**
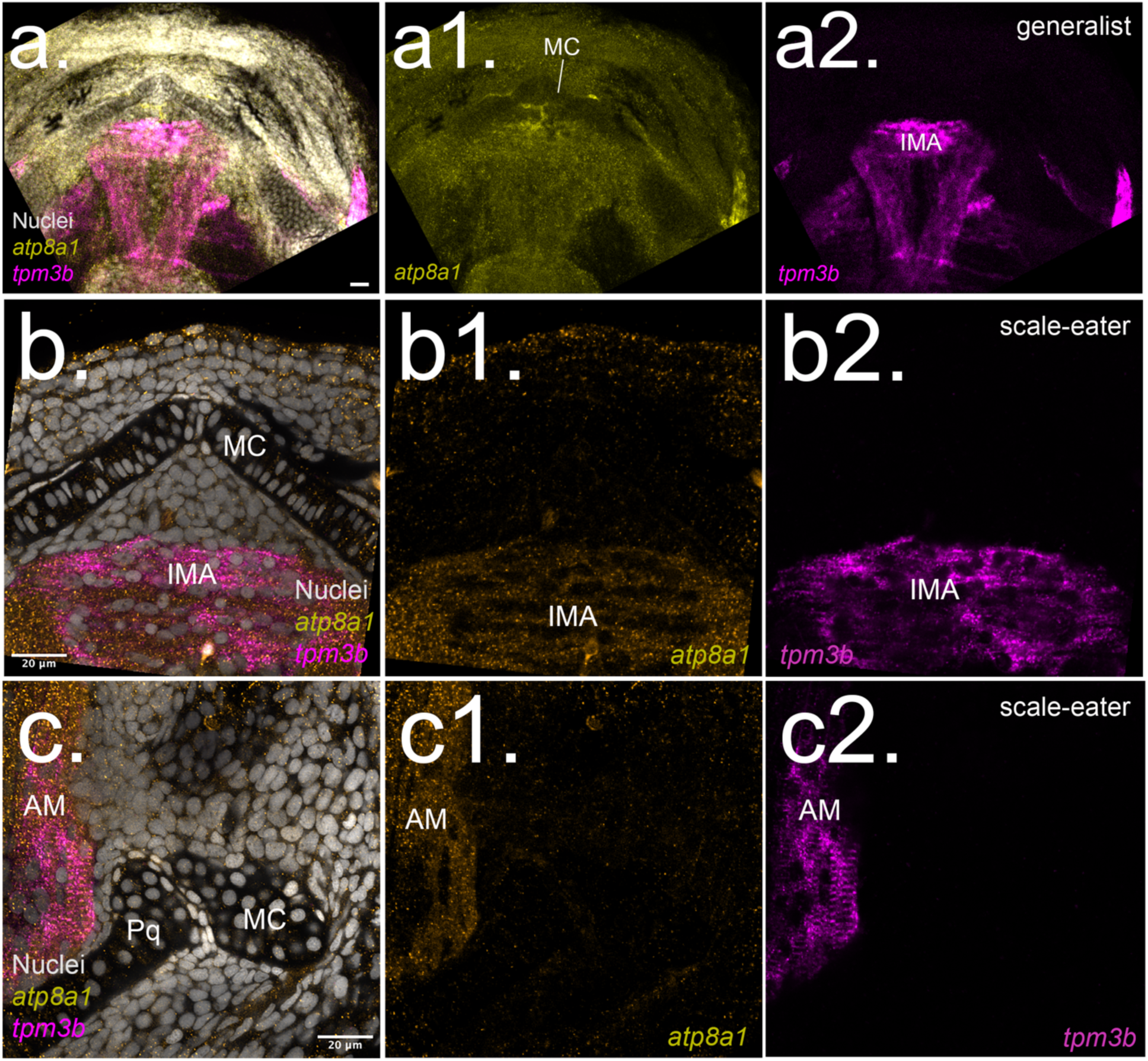
*Atp8a1* expression is restricted to the developing intermandibular anterior and adductor mandibulae craniofacial muscles in scale-eaters. Single optical section of fluorescent hybridization chain reaction (HCR) for *atp8a1* (orange) and *tpm3b* (magenta) in the lower jaw of a) generalist and b-c) scale-eaters at hatching (8 dpf). a1-c1) Single channel for *atp8a1*. a2-c2) Single channel for *tpm3b*. MC: Meckel’s cartilage, IMA: intermandibular anterior muscle, AM: adductor mandibulae muscle. Nuclei were labeled with DAPI.

## Discussion

Understanding how genetic variants regulate gene expression and drive morphological variation among species is fundamental for understanding the genetic basis of both human diseases and adaptive evolution. In this study, we used tissue-specific RNA sequencing within a nascent adaptive radiation of pupfishes on San Salvador Island to investigate the genetic changes underlying differential gene expression during development among these highly divergent trophic specialists. From over 5 million segregating single nucleotide polymorphisms and thousands of loci previously associated with oral jaw divergence in these species (Richards et al. 2021, 2022, McGirr and Martin 2021), we pinpointed a tiny subset (*n* = 9) that overlap with differentially expressed genes (DEGs) unique to craniofacial tissues in pupfish trophic specialists at hatching (8 dpf).

Furthermore, we additionally validated divergent and novel craniofacial expression in two of these genes using in situ hybridization, focusing on the two most prominent candidates based on previous genomic studies in this system. Notable, all three candidate genes investigated thus far - including *galr2* from an earlier work- have revealed novel expression patterns and putative roles in vertebrate craniofacial development, representing an unexpected success rate of 100%. This demonstrates the power of the pupfish system to discover novel gene functions and sheds light on the regulatory mechanisms shaping craniofacial in trophic specialists. Moreover, our findings highlight the utility of integrating tissue-specific transcriptomics with evolutionary genomics to reveal new insights into the genetic architecture of adaptive morphological divergence.

### Increased craniofacial transcriptional variance reflects exceptional morphological diversification in this region

We observed significantly higher transcriptional variance in craniofacial tissues (Fig. 2 c, PC1: 48.6%) than in caudal region tissues (Fig. 2e), a pattern that was consistent even when limited to San Salvador Island species (PC1: 61.3% vs. PC1: 22.9% in craniofacial versus caudal tissues, respectively). This elevated variance aligns with the exceptionally fast rates of morphological diversification observed primarily in craniofacial traits, while the caudal region has remained relatively conserved across all *Cyprinodon* species (Martin 2016; Martin and Wainwright 2011). Interestingly, the largest effect sizes for the 30 trait-associated QTL examined to date have also been identified only in craniofacial traits, accounting up to 15% of oral jaw size variation, but not for caudal traits (Martin, Erickson, and Miller 2017). Our findings suggest that rapid morphological diversification may be linked to shifts in the underlying genetic architecture (also see (Chan et al. 2024; St John et al. 2024)), and here we show an additional association with elevated transcriptional variance.

### Sensory perception as an unexplored factor for trophic specialization and craniofacial divergence in Bahamian pupfishes

Within the craniofacial-exclusive DEGs shared among trophic specialists, we observed significant enrichment for biological processes related to ‘sensory perception’ (Fig. 4, b-c) and ‘response to stimulus’ (Fig. 4, b-c). Most of the DEGs shared among specialists were associated with ‘visual perception’ and ‘signal transduction in cell from the retina’, and expressed in the visual system. Genes such as *six3b, neurod1,* and *neurod4* are known to be involved in nervous system development and neuronal differentiation, with *six3b* playing a role in the development of the visual placode and *neurod4* to the olfactory placode. This raises an intriguing question: why are these 71 craniofacial-exclusive genes shared among trophic specialists yet differentially expressed relative to other pupfish species?

The sensory drive theory of evolution predicts that variation in sensory perception among different populations, driven by divergent ecological selection, can promote speciation by influencing mating traits and reproductive isolation (Endler 1992, 1993; Boughman 2002; Cummings, Endler, and Handling 2018). The enrichment of sensory processing genes among the specialist-shared craniofacial-exclusive DEGs (Fig. 4, b, c; Table S4), coupled with previous findings from our lab linking highly differentiated adaptive variants to sensory development genes (Richards and Martin 2022), suggest that sensory perception has played a critical role in pupfish speciation on SSI. Growing evidence across taxa -particularly in fish- points to the role of environmental tuning favoring vision over other sensory modalities during speciation especially in fishes (Kawata et al. 2007; Seehausen et al. 2008; Rennison et al. 2016; Marques et al. 2017; Cummings, Endler, and Handling 2018; Abate and Noakes 2021), supporting the idea that eye development and retinal signal transduction may have been crucial in the evolution of trophic specialization in SSI pupfishes.

### Craniofacial expression of pycr3 highlights a role for proline metabolism in the evolution of scale-eating morphological divergence

Unlike Pycr1 and Pycr2, which are found in mitochondria, the pyrroline-5-carboxylate reductase Pycr3 localizes in the cytosol, where it specifically catalyzes the synthesis of proline from ornithine-derived Δ1-Pyrroline-5-carboxylate (P5C). Besides their role in proline metabolism, Pycr3, as well as the other pyrroline-5-carboxylate reductases Pycr1 and Pycr2, also plays a role in controlling dermal elasticity, oxidative stress response, cancer cell proliferation, and preventing aging and apoptosis (Beyens et al. 2021; Reversade et al. 2009; Liang et al. 2019; Kuo et al. 2016). Our study demonstrates that *pycr3* is differentially expressed in the craniofacial tissue of *C. desquamator* scale-eaters in comparison with all other pupfish species at hatching. Specifically, *pycr3* is expressed in cranial neural crest (CNC) derivatives, including scattered cells throughout the brain (Fig. 6, a-d), the mesenchyme surrounding the future dentary and premaxilla symphyses (Fig. 6, c-d), the chondrocranium (Fig. 6, c), and the gill arches of generalist pupfishes, but absent in scale-eater gill arches (Fig. 6, a-b). A recent zebrafish *pycr3* knock-out model (*Identification Characterization Genetic Variation Zebrafish Hsp90ab1 Locus Reveal Novel Role Pycr3 Neural Crest Cell Biology Stress Response*, unpublished thesis) revealed abnormalities in skeletal structures and CNC-derivatives in embryos and larvae, including spinal curvature, abnormal jaw size, and a reduced head and eye size. Although proline metabolism is unaffected in *pycr3^-/-^* zebrafish embryos there is a reduction in glutathione levels suggesting that proline metabolism – but not redox transfer regulation- is being compensated by other effectors, such as Pycr1 and/or Pycr2 (*Identification Characterization Genetic Variation Zebrafish Hsp90ab1 Locus Reveal Novel Role Pycr3 Neural Crest Cell Biology Stress Response*, unpublished thesis). We further analyze the expression of the annotated *pycr1a* and *pycr1* gene products in our dataset and found them to be expressed at similar levels across species and tissues (Fig. S2), supporting a specific role for *pycr3* in the divergent craniofacial development of scale-eater pupfish

Our work therefore extends the understanding of *pycr3*’s novel role in fish craniofacial development while also supporting a specific evolutionary contribution of *pycr3* and proline metabolism to the development of jaw morphological divergence in scale-eating pupfishes. Whether the absence or near absence of *pycr3* expression in the craniofacial tissue of *C. desquamator* and ‘wide-mouth’ scale-eaters at hatching, compared to generalists and the molluscivores, respectively (Fig. 5, e1; 6) is (i) a consequence of heterochrony (Holtmeier 2001) and/or (ii) a direct result of cis or trans regulation of gene expression (McGirr and Martin 2021) remains to be elucidated.

### Novel differential expression of atp8a1 supports a role for muscle development in craniofacial divergence of scale-eating pupfish

Atp8a1 is member of the P4-ATPase flippase type A family complex that acts in association with CDC50a proteins, catalyzing the coupled hydrolysis of ATP and the translocation of phosphatidylserine (PS) to the inner leaflet of the membrane, thereby altering the PS distribution to facilitate plasma membrane vesicle formation across taxa (Kook et al. 2021; Hasegawa et al. 2021; Best, Xu, and Graham 2019). Intracellularly, Atp8a1 localizes to the trans-Golgi network, endoplasmic reticulum, endosomes, and lysosome-related organelles (Pocognoni et al. 2024; Kook et al. 2021). In mammals, Atp8a1 is broadly-expressed across tissues, including the central nervous system, retina, spleen, bladder, testes, kidney, lung, heart, and skeletal muscles (Wang et al. 2018). Although the physiological role of P4 ATPases remain poorly understood, reported mutations in humans have been linked to various chronic and progressive liver diseases (Panatala, Hennrich, and Holthuis 2015). In mice, an Atp8a1 knockout model is non-lethal and displays no apparent morphological abnormalities apart from disrupted hematopoietic stem cell pool homeostasis (Zheng et al. 2023) and behavioral hyperactivity with impaired hippocampus-associated learning (Levano et al. 2012).

The function and expression of Atp8a1 in fish, prior to this work, remained largely unexplored. Similar to the expression other P4-ATPases in mammals, *atp11a, atp11b,* and *atp11c,* have shown expression in the zebrafish neural crest cells and some of their derivatives, including the pharyngeal arches, the inner ear, and the retina (Hawkey-Noble et al. 2020). Tissue-specific transcriptomics showed a reduced expression of *atp8a1* in both SSI scale-eater pupfishes, significantly different for the newly discovered intermediate *C.* ‘wide-mouth’ (Fig. 5, f). The restricted expression of *atp8a1* to *tpm3b*-positive craniofacial muscles, such as the AM and IMA, in *C. desquamator* scale-eaters (Fig. 7, b-c) suggests that *atp8a1* expression is subject to different spatially and/or temporally regulation among scale-eaters compared to generalists. In mice, the usually observed high expression of *atp8a1* in adult skeletal muscle (Tsuchiya et al. 2018) shows decreases associated with reduced chromatin accessibility of its regulatory regions upon denervation, exemplifying how *atp8a1* transcriptional regulation occurs in an atrophy-related responsive circuit (Lin et al. 2023). Our findings reveal, for the first time, a differential expression pattern of *atp8a1* in the craniofacial muscles of any vertebrate, indicating a novel role in the evolution of craniofacial morphogenesis. However, how *atp8a1* contributes to differential craniofacial morphogenesis between pupfish species remains unknown, ensuring future directions exploring the role of *at8a1* in craniofacial muscle evolution and development.

Across vertebrates, during craniofacial development, neural crest cells and the surface ectoderm interact with the adjacent pharyngeal mesoderm within a recently described developmental organizer domain, the cardiopharyngeal field, directing its differentiation into either cardiac or craniofacial skeletal muscle (Nathan et al. 2008; Diogo et al. 2015; Michailovici, Eigler, and Tzahor 2015). It remains to be determined whether *pycr3* and *atp8a1* are expressed in neural crest cells or the pharyngeal mesoderm during early pupfish embryogenesis, the specific embryonic stage at which *pycr3* expression becomes evident in the gill arches of molluscivores, and when *atp8a1* expression becomes restricted to the *tpm3b*-positive craniofacial muscles in scale-eaters. Given their expression at hatching stage, we hypothesize that *pycr3* may be found expressed in cranial neural crest cells of generalists and molluscivores at the pharyngula stage, whereas *atp8a1* may be expressed in the pharyngeal mesoderm, as it later shows expression in the craniofacial muscles of pupfish hatchlings (this study) and in both developing and adult mammalian skeletal muscle (Gene Expression Omnibus (GEO) [Internet]. Bethesda (MD): National Center for Biotechnology Information (US); 1999). Whether these differential gene expression changes influence behavior, or whether the fixed adaptative alleles identified in the specialist species directly regulate the specialist-specific and craniofacial-exclusive genes reported here (Fig. 5, a-c) via genetic regulatory mechanisms (i.e., cis or trans regulation), as well as the impact of these regulatory pathways on skeletal heterochrony during craniofacial development between pupfishes, remains to be studied.

## Acknowledgments

This research was funded by NSF CAREER 1749764 and NIH 5R01DE027052-02 to CHM. We thank the Martin lab for their valuable comments and discussion of the results and the manuscript. We also thank Lydia Smith for her help in the Evolutionary Genetics Lab at the University of California, Berkeley; the Gerace Research Center and Troy Day for logistical support in the field, and the government of the Bahamas for permission to collect and export samples in 2018. All research procedures and animal care protocols (AUP-2021-02-14062-1 and AUP-2021-07-14515) were approved by the University of California, Berkeley Animal Care and Use committee.

## Data Accessibility

All sequencing reads generated by this study will be deposited in the NCBI Sequence Read Archive once the paper is published. Scripts for analysis and visualization of the results are available on the author’s GitHub, github.com/fishfena/JAWesome_RNAproject.

## Author Contributions

M.F.P.: conceptualization, formal analysis, investigation, visualization, writing-original draft, writing-review & editing; V.M: investigation; C.H.M: conceptualization, funding acquisition, resources, writing-review & editing, validation.

## Notes

### Competing Interest Statement

The authors have declared no competing interest.

