## Supplemental Figures and Tables for "Tissue-specific transcriptomics uncovers novel craniofacial genes underlying jaw divergence in specialist pupfishes"

Figure S1- S2, Pages 1 - 2.

**Supplemental Tables**

Tables S1-S5, Pages 3-7.

### Supplemental Figures

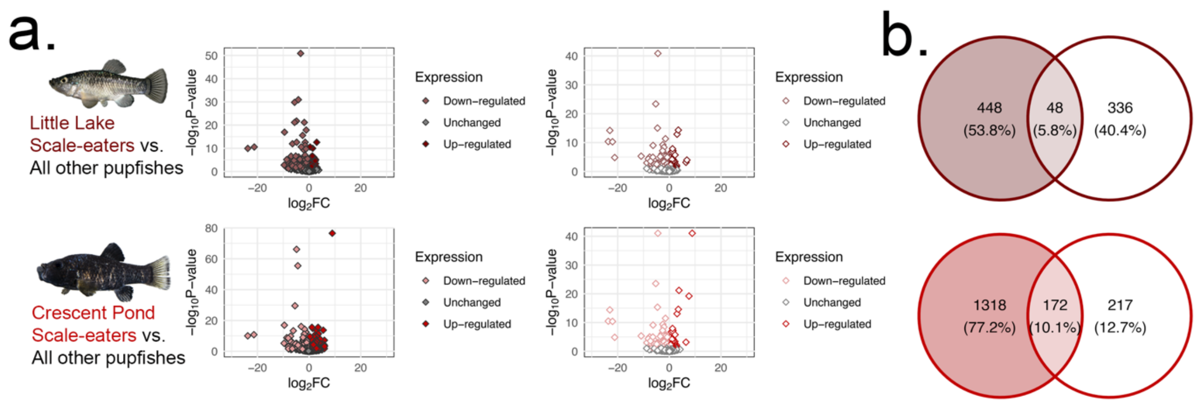


**Figure S1. Differential gene expression in two different lake populations of *C. desquamator* relative to all other pupfishes.** Craniofacial tissue (column 1: filled symbols) and caudal region tissue (column 2: open symbols) for Little Lake (first row) and Crescent Pond (second row). a) Volcano plots showing up and downregulated genes. b) The number of tissue- and population-exclusive differentially expressed genes.

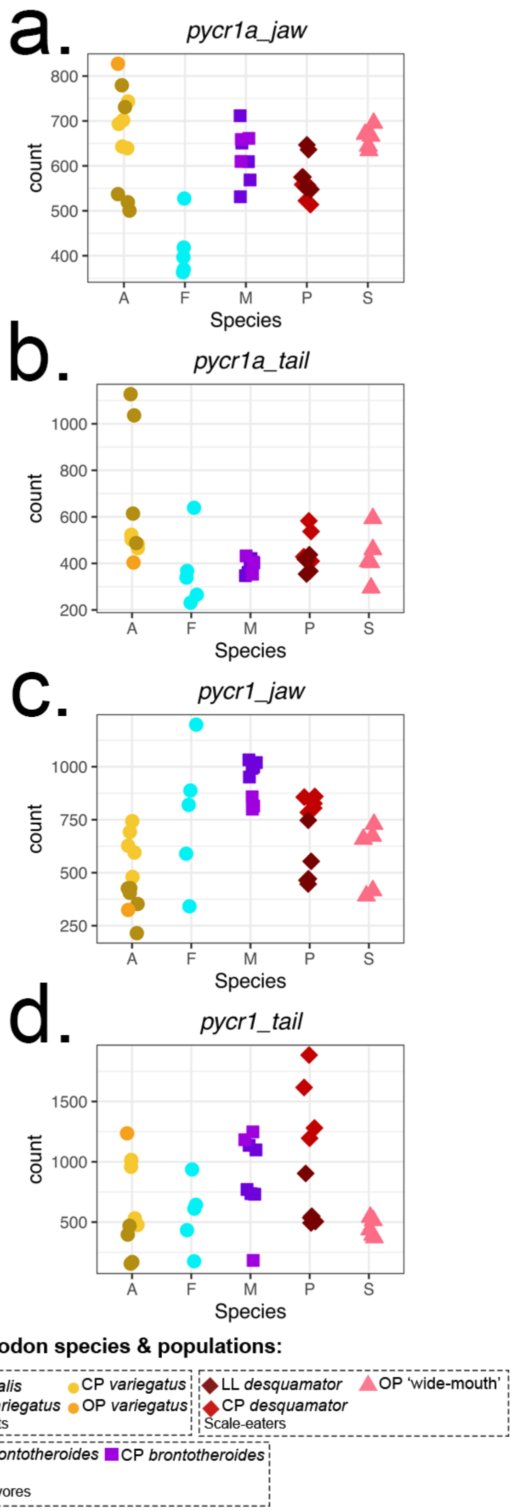


**Figure S2. *Pycr1a* and *pycr1* expression across tissues and populations.** Pycr1a is encoded by ENSCVAG00000011390 and *pycr1* by ENSCVAG00000019911.

### Supplemental Tables

**Table S1.** Number of replicates of each sequences RNA sample. Populations are abbreviated as, CRP: Crescent Pond, OSP: Osprey Lake, NC: North Carolina, LIL: Little Lake.

| ***Cyprinodon*** | **Population** | **Tissue** | **# *Replicates*** |
|---|---|---|---|
| *variegatus* | CRP | Jaw | 5 |
| *variegatus* | OSP | Jaw | 1 |
| *variegatus* | NC | Jaw | 5 |
| *brontotheroides* | CRP | Jaw | 5 |
| *brontotheroides* | OSP | Jaw | 3 |
| *desquamator* | CRP | Jaw | 5 |
| *desquamator* | LIL | Jaw | 5 |
| *wide-mouth* | OSP | Jaw | 5 |
| *fontinalis* | | Jaw | 5 |
| *variegatus* | CRP | Jaw | 4 |
| *variegatus* | OSP | Jaw | 1 |
| *brontotheroides* | CRP | Jaw | 4 |
| *brontotheroides* | OSP | Jaw | 5 |
| *desquamator* | CRP | Jaw | 3 |
| *desquamator* | LIL | Jaw | 4 |
| *wide-mouth* | OSP | Jaw | 5 |
| *fontinalis* | | Jaw | 5 |

**Table S2.** Three adaptive genes are differentially expressed between molluscivores and all other pupfishes at hatching. $P < 0.01$ in bold. LFC, log2 fold change in expression (negative values indicate lower expression in molluscivores than other pupfishes); MNC, mean normalized read counts across samples; $P$, adjusted $P$ values for differential expression (DESeq2).

| **Gene** | **MNC** | **LFC** | ***P*** |
|---|---|---|---|
| *coq7* | 60.40 | -1.51 | **1.92E-04** |
| *gga1* | 143.12 | -0.50 | 2.38E-02 |
| *mylipa* | 648.98 | -0.79 | 1.08E-02 |

**Table S3.** Five adaptive genes are differentially expressed between scale-eaters and all other pupfishes at hatching. *P* < 0.01 in bold. LFC, log2 fold change in expression (positive values indicate higher expression in the scale-eaters than in other pupfishes while negative values indicate lower expression); MNC, mean normalized read counts across samples; *P*, adjusted *P* values for differential expression (DESeq2).

| **Gene** | **MNC** | **LFC** | ***P*** |
|---|---|---|---|
| *pycr3* | 10.11 | -6.81 | **3.78E-13** |
| *slc51a* | 106.99 | 0.47 | 4.50E-02 |
| *bri3bp* | 147.95 | 0.43 | 1.13E-02 |
| *vgll3* | 141.60 | -0.47 | **2.67E-03** |
| *mag* | 658.46 | -0.42 | 4.33E-02 |

**Table S4.** One adaptive gene is differentially expressed between the intermediate scale-eater *C.* 'Wide-mouth' and all other pupfishes at hatching. LFC, log2 fold change in expression (negative values indicate lower expression in intermediate scale-eaters than other pupfishes); MNC, mean normalized read counts across samples; *P*, adjusted *P* values for differential expression (DESeq2).

| **Gene** | **MNC** | **LFC** | ***P*** |
|---|---|---|---|
| *atp8a1* | 773.33 | -0.34 | 4.43E-02 |

**Table S5.** Gene Ontology functional categories (using ShinyGo) for craniofacial-exclusive differentially expressed genes shared between specialists. Categories with significant false discovery rate (FDR) are bolded. Genes without annotations are listed as Ensemble transcripts.

| **GO category** | **Genes** |
|---|---|
| Regulation of cellular process | *grk1a, opn1sw2, grm6b, six3b, ccny, opn4xb, rgrb, neurod1, ENSCVAT00000017314, rgs7bpb, gngt1, ENSCVAT00000030681, prdm1b, sagb, ccdc88aa* |
| **Response to stimulus** | *grk1a, opn1sw2, grm6b, rcvrna, opn4xb, rgrb, neurod1 ENSCVAT00000017314, rgs7bpb, gngt1, ENSCVAT00000030681, sagb, cnga3* |

| | |
|---|---|
| Signaling | *grk1a, opn1sw2, grmb6b, sv2ba, opn4xb, rgrb, neurod1, ENSCVAT00000017314, rgs7bpb, gngt1, ENSCVAT00000030681, sagb* |
| Localization | *sv2ba, vtg5, abca4a, ENSCVAT00000010196, aqb9b, ENSCVAT00000003747, ENSCVAT00000016798, slc1a7a, cplx4b, ccdc88aa, cnga3, ENSCVAT00000028392* |
| Establishment of localization | *sv2ba, vtg5, abca4a, ENSCVAT00000010196, aqb9b, ENSCVAT00000003747, ENSCVAT00000016798, slc1a7a, cplx4b, ccdc88aa, cnga3, ENSCVAT00000028392* |
| **Cellular response to stimulus** | *grk1a, opn1sw2, grmb6b, opn4xb, rgrb, neurod1, ENSCVAT00000017314, rgs7bpb, gngt1, ENSCVAT00000030681, sagb* |
| Multicellular organismal process | *neurod4, grk1a, opn1sw2, grmb6b, ENSCVAT00000011047, six3b, rgrb, neurod1 ENSCVAT00000016798, impg2a* |
| System process | *grk1a, opn1sw2, grmb6b, ENSCVAT00000011047, rgrb, ENSCVAT00000016798, impg2a* |
| Developmental process | *neurod4, ENSCVAT00000011047, six3b, neurod1 impg2a, prdm1b* |
| **Response to abiotic stimulus** | *grk1a, opn1sw2, grmb6b, ENSCVAT00000011510, opn4xb, rgrb* |
| **Response to external stimulus** | *grk1a, opn1sw2, grmb6b, opn4xb, rgrb* |
| Anatomical structure development | *neurod4, ENSCVAT00000011047, six3b, neurod1 impg2a,* |
| **Detection of stimulus** | *grk1a, opn1sw2, grmb6b, opn4xb, rgrb,* |
| Cellular component organization or biogenesis | *ino80c, pmelb, ccdc88aa, tmem138* |
| Biosynthetic process | *six3b, neurod1, ENSCVAT00000017314, prdm1b* |
| Cellular component organization | *ino80c, pmelb, ccdc88aa, tmem138* |
| Regulation of metabolic process | *six3b, ccny, neurod1 prdm1b* |
| Negative regulation of biological process | *neurod1 rgs7bpb, prdm1b* |
| Regulation of signaling | *grk1a, neurod1, rgs7bpb* |
| **Regulation of response to stimulus** | *grk1a, neurod1, rgs7bpb* |
| Regulation of molecular function | *ENSCVAT00000008171, ccny, ENSCVAT00000000410* |
| Cell adhesion | *ENSCVAT00000010679, pcdh8* |
| Anatomical structure morphogenesis | *six3b, neurod1* |
| Negative regulation of signaling | *neurod1, rgs7bpb* |

| | |
|---|---|
| Methylation | *prdm1b, prdm13* |
| Growth | *six3b* |
| Locomotion | *ccdc88aa* |
| Catabolic process | *eno1b* |
| Macromolecule localization | *vtg5* |
| Hormone metabolic process | *ENSCVAT00000017446* |
| Pigmentation | *pmelb* |
| Cellular component biogenesis | *tmem138* |
| Developmental growth | *six3b* |
| Cell motility | *ccdc88aa* |
| Cellular localization | *ccdc88aa* |
| Localization of cell | *ccdc88aa* |
| Regulation of biological quality | *ENSCVAT00000017446* |
